# Visually evoked responses are enhanced when engaging in a video game

**DOI:** 10.1101/588806

**Authors:** Jason J. Ki, Lucas C. Parra, Jacek P. Dmochowski

**Affiliations:** Department of Biomedical Engineering, City College of New York, New York NY 10031

## Abstract

While it is well known that vision guides movement, less appreciated is that the motor cortex also provides input to the visual system. Here we asked whether neural processing of visual stimuli is acutely modulated during motor activity, hypothesizing that visual evoked responses are enhanced when engaged in a motor task that depends on the visual stimulus. To test this, we told participants that their brain activity was controlling a video game that was in fact the playback of a prerecorded game. The deception, which was effective in half of participants, aimed to engage the motor system while avoiding evoked responses related to actual movement or somatosensation. In other trials, subjects actively played the game with keyboard control or passively watched a playback. The strength of visually evoked responses was measured as the temporal correlation between the continuous stimulus and the evoked potentials on the scalp. We found reduced correlation during passive viewing, but no difference between active and sham play. Alpha band (8-12 Hz) activity was reduced over central electrodes during sham play, indicating recruitment of motor cortex despite the absence of overt movement. To account for the potential increase of attention during game play, we conducted a second study with subjects counting screen items during viewing. We again found increased correlation during sham play, but no difference between counting and passive viewing. While we cannot fully rule out the involvement of attention, our findings do demonstrate an enhancement of visual evoked responses during active vision.

## Introduction

Visual processing in the brain has historically been delineated into two streams corresponding to the primary roles of vision: perception and action (1, 2). The visual guidance of actions manifests as a sequence of transformations along the so-called “dorsal visual stream”, originating in the primary visual cortex and passing through the parietal lobe (3, 4) before terminating in the motor cortex. Studies of visually-guided action have generally adopted a feedforward view, where the relevant information flows from the visual system to the premotor and motor centers. On the other hand, much less attention has been devoted to potential influences of downstream regions, including the motor cortex itself, on visual processing.

Despite this, multiple lines of evidence indicate that the motor system exerts influence over visual processing. First, the visual and motor cortices have reciprocal anatomical connections in the primate brain (5–7). Moreover, numerous behavioral studies have demonstrated that learning of motor actions improves subsequent recognition of congruent visual stimuli (8–10), and that perceptual decisions may be primed by action (11–13). Human sensitivity to visual motion appears to be higher when that motion matches the observer’s own movement patterns (14, 15). There is also evidence from neuroimaging studies that objects affording actions enhance early visual evoked potentials (VEPs) via a purported sensory gain mechanism (16–19). Neural recordings from visual extinction patients demonstrate that graspable objects bias visual perception in an unconscious manner (20). Peripheral visual processing in the human brain is enhanced during locomotion (21). Based on these findings, we suspected that the presence of a motor task could *acutely* modulate visual processing. Specifically, we hypothesized that visual evoked responses are enhanced when the stimulus guides motor control. Testing this hypothesis in the human brain is not straightforward due to the fact that manual actions (e.g. button presses) introduce somatosensory and motor related signals into recordings of brain activity, potentially confounding measures of neural visual responses, particularly because actions are often time-locked to changes in the stimulus.

Here we developed a “sham” motor task aimed at identifying the online effect of motor engagement on the dynamics of concurrent visual processing (throughout the paper, “motor engagement” implies that the motor activity is coupled with the visual stimulus). Subjects were under the belief that their brain activity was controlling a car racing video game, when in fact they were viewing a recording. The purpose of this manipulation was to engage the motor cortex while avoiding somatosensory or motor evoked potentials. Neural activity was recorded with the scalp electroencephalogram (EEG) to capture fast neural responses that could then be correlated with rapid stimulus fluctuations without requiring exogenous stimulus labels. We assessed the strength of visual evoked responses by measuring the correlation between a feature of the time-varying visual stimulus (optic flow) and the evoked EEG response: the stimulus-response correlation (SRC) (22, 23). To test the hypothesis of enhanced visual evoked responses during motor engagement, we compared the SRC obtained during “sham play” with that measured during passive viewing and conventional manual game play (“active play”). To investigate the possible role of increased attention during video game play, we conducted a second study where subjects were additionally asked to view the game while counting the appearance of target items on the screen. In what follows, we report a robust enhancement of visual evoked responses when subjects were engaged in video game play, and discuss the roles of motor coupling and attention in our findings.

## Results

We hypothesized that visual evoked responses are enhanced during stimulus-dependent motor control. To test our hypothesis while ruling out activity associated with actual movement, we informed study participants that their brain activity would be controlling a car racing video game but instead presented them with playback of a previously recorded game (“sham play”). In other trials, subjects controlled the game with keyboard presses (“active play”) or passively viewed game playback (“passive viewing”; Fig 1A).

**Fig. 1.**
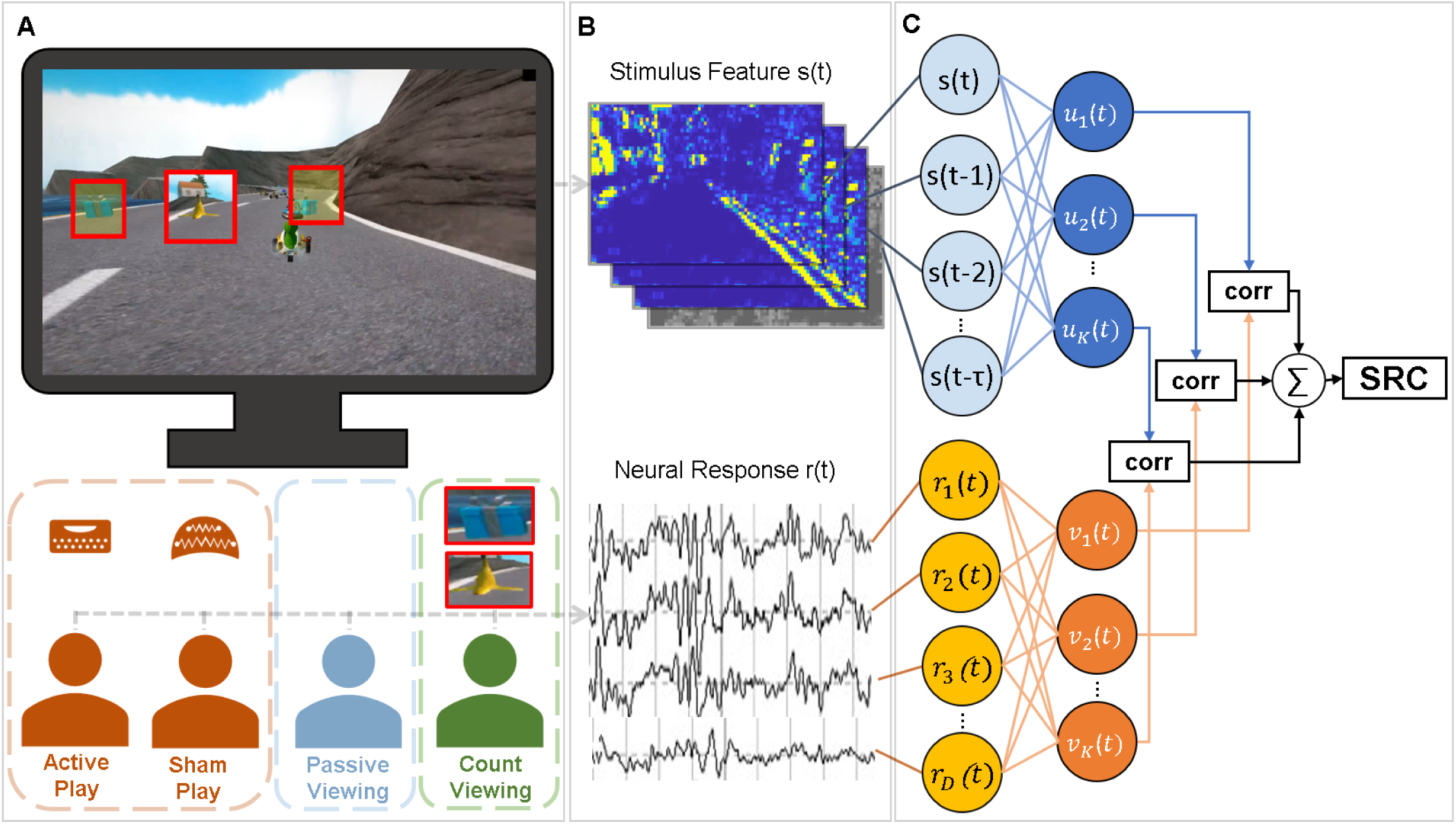
Measuring visual evoked responses with and without motor engagement. (**A**) Study participants experienced a car racing video game under several conditions: manual control (“active play”), viewing but under the false belief that brain activity was controlling game play (“sham play”), and knowingly viewing game playback (“passive viewing”). In a follow up study, we added an attention demanding task (“count viewing”) to control for the possible effects of top-down attention. (**B**) Throughout the experiment, we recorded the video stream as well as the evoked scalp EEG. (**C**) The strength of visual evoked responses was assessed by measuring the temporal correlation between the overall optic flow of the video stream and the time-locked neural response. To account for varying response latencies and multiple recording electrodes, we formed multiple spatial components of the EEG and temporal components of the stimulus using Canonical Correlation Analysis (22, 23). The sum of correlations across all components formed the dependent measure, which we term here the stimulus-response correlation (SRC).

Our dependent measure was the temporal correlation between the time-varying optic flow of the video stream and the evoked brain response captured by the scalp EEG (Fig 1B). To account for the spatial diversity of the 96-channel EEG and varying response latencies, we captured multiple spatial components of the EEG and temporal components of the stimulus following the methodology developed previously (22, 23). This approach employs Canonical Correlation Analysis (CCA) to model neural responses to continuous stimuli with “temporal response functions” (24). These evoked responses are entirely equivalent to conventional event-related potentials (ERP) but do not require the specification of discrete visual events. The CCA approach differs from multivariate regression in that it decomposes neural activity into components with their own temporal and spatial profile. We measure the overall strength of the visual evoked responses as the summed correlation measured in each component to arrive at the total stimulus-response correlation (SRC; Fig 1C).

When applied to the present data, we obtained several visual response components evoked by optic-flow fluctuations (Fig 2A). Notably, the strongest component was marked by a parietal topography centered at electrode CPz (centroparietal midline). The corresponding temporal response function showed a positive peak at 200 ms. The second strongest component exhibited poles over the medial frontal and medial occipital regions, and showed a late temporal response with a peak at 400 ms (Fig 2A). Components 3 and 4 showed mirror symmetric spatial response functions with peak expression over right and left frontocentral electrodes, respectively. Collectively, the set of evoked response functions indicate that the visual stimulus drove neural activity over broad scalp regions and included late responses.

**Fig. 2.**
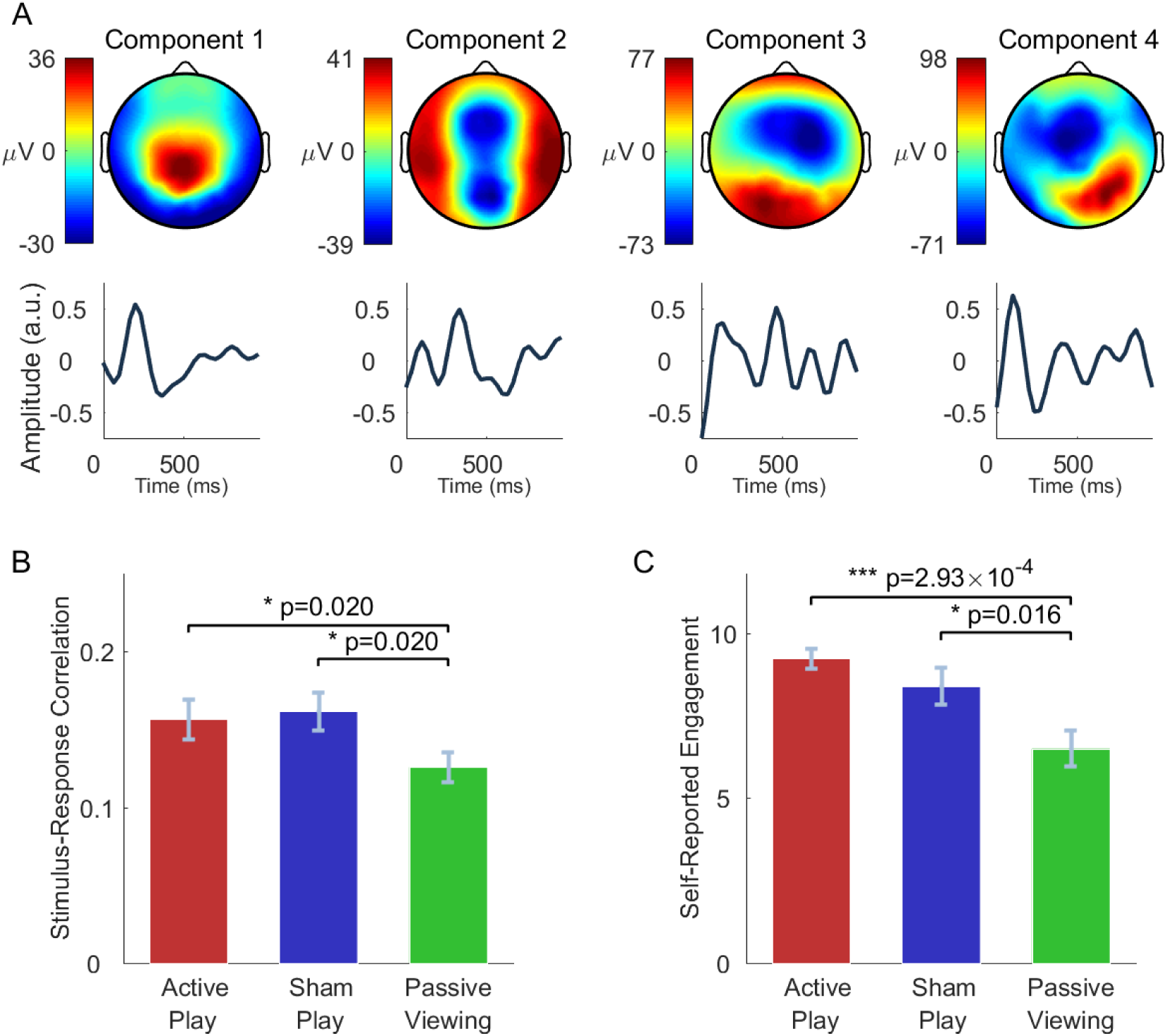
Enhanced visual evoked responses during active and sham play. (**A**) Spatial and temporal response functions for the four strongest components evoked by the optic flow of the video game stimulus. Time indicates the delay of the EEG evoked response relative to the stimulus presentation time. (**B**) The total SRC was measured separately for each condition (bar height depicts mean ± sem across *n* = 18 subjects). Passive viewing elicited significantly lower total SRC compared to active play (*p* = 0.02, *n* = 18, paired two-tailed Wilcoxon signed rank test) and compared to sham play (*p* = 0.02, *n* = 18). No significant difference was found between active and sham play (*p* = 0.58, *n* = 18). (**C**) Participants were asked to rate their engagement with the video game in each condition. Subjects reported significantly higher engagement during active play (*p* = 2.93 × 10^−4^, *n* = 18, paired two-tailed Wilcoxon signed rank test) and sham play (*p* = 0.016, *n* = 18) relative to passive viewing. No significant difference in self-reported engagement was found between active and sham play (*p* = 0.24, *n* = 18).

### Enhanced stimulus-response correlation during active and sham play

We measured the total SRC separately for each experimental condition and found a significant increase during sham play relative to passive viewing (*z* = 2.33, *p* = 0.02; paired, two-tailed Wilcoxon signed-rank test, *n* = 18 subjects; Fig 2B). Similarly, SRC was increased during active play (*z* = −2.33, *p* = 0.02, Fig 2B). No significant difference in SRC was found between active and sham play (*z* = −0.54, *p* = 0.58; Fig 2B). To compute the SRC, we employed the optic flow of the video stream because this particular feature drives the EEG stronger than other low-level visual or auditory features (22). However, similar results were obtained with temporal visual contrast (Figure S1). Namely, the spatial response functions are highly congruent (compare Figs S1A and 2A), and we found a significant increase in SRC during sham play compared to passive viewing (*z* = 2.61, *p* = 0.008, Fig S1B), and a numerically higher SRC during active play relative to passive viewing (*z* = 1.48, *p* = 0.138). We therefore continued our analysis with the optic flow feature.

Following the experiment, participants were asked to rate their engagement with the game in each condition. Analogous to the SRC measure, subjects reported higher engagement scores for active play (*z* = 3.44, *p* = 2.9 · 10^−4^, paired, two-tailed Wilcoxon signed-rank test, *n* = 18 subjects) and sham play (*z* = 2.13, *p* = 0.031) relative to passive viewing (Fig 2C). No significant difference in self-reported engagement was observed between active and sham play (*z* = 1.17, *p* = 0.24; Fig 2C).

After completing the post-experiment survey, subjects were informed of the deception in the sham play, and were asked whether they had become aware of the fact that their brain activity was not controlling game play. Of the 18 study participants, 13 reported being deceived for the entirety of the experiment. The remaining five subjects did not immediately notice the sham. Since this evaluation is subjective and all participants may have been deceived for at least some of the time, we decided to not make a distinction between the two groups of subjects during data analysis.

### Alpha desynchronization over motor cortex indicates motor engagement during sham play

By design, there were no overt differences in behavior between sham play and passive viewing – in both conditions, participants viewed the stimulus without performing manual actions. This prevented confounds due to motor or somatosensory evoked responses that could have been present during active play. To test whether our sham condition nevertheless engaged motor cortex, we measured the power of alpha-band (8-12 Hz) oscillations for each condition. Desynchronization of alpha activity has long been observed over the motor cortex (“mu” rhythm) when subjects perform or visualize motor actions (25). Indeed, we observed a significant reduction in alpha power during both active and sham play relative to passive viewing, with the largest differences observed at bilateral central scalp locations over the motor cortex (Fig 3A-B). On the other hand, alpha power did not significantly differ between active and sham play (Fig 3B). This provides evidence that the motor system was indeed engaged during sham play.

**Fig. 3.**
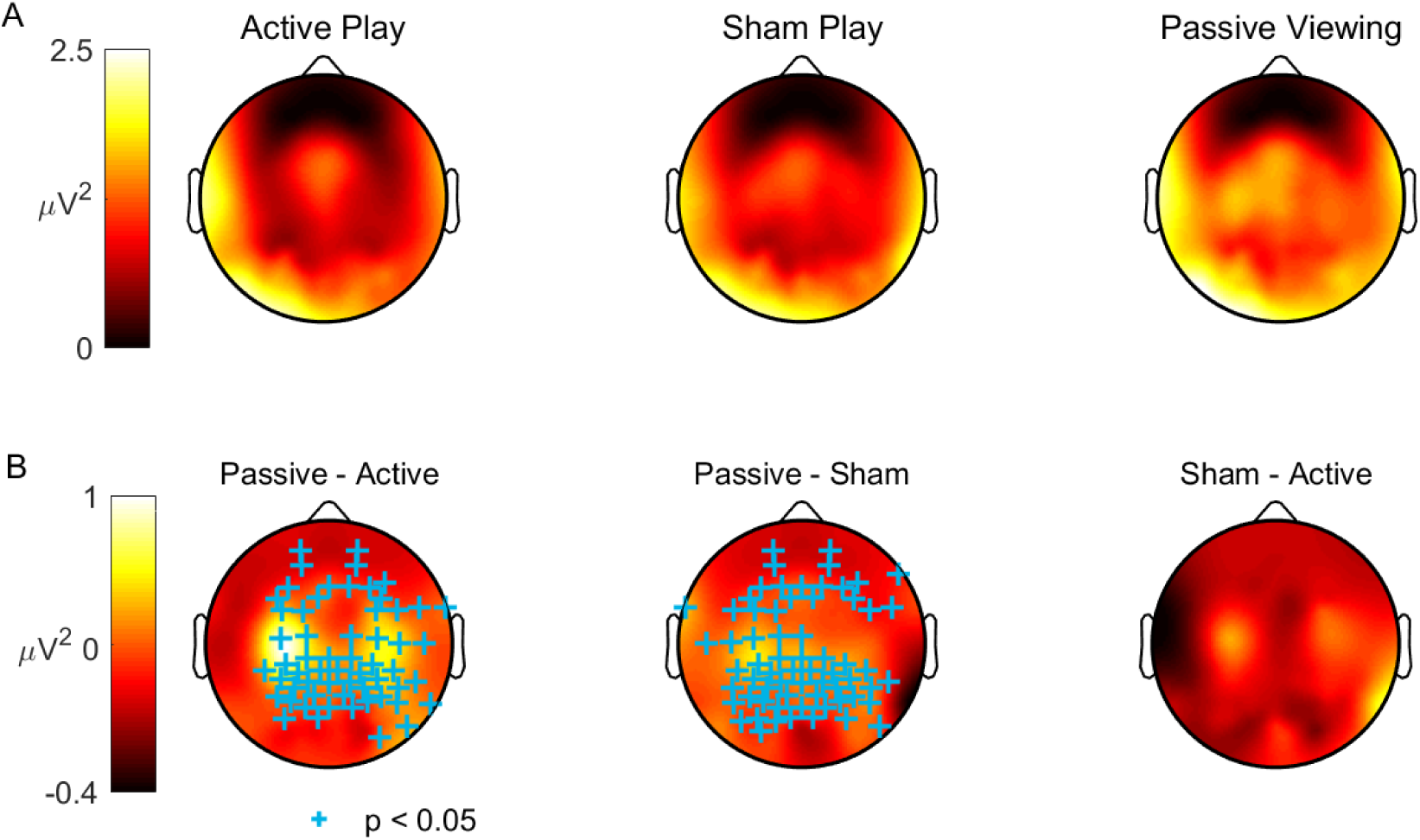
Alpha desynchronization over motor cortex during active and sham play. (**A**) The power of the EEG in the alpha band (8-12 Hz) across the scalp, shown for each experimental condition. Note the greater power over left central locations during passive viewing. (**B**) The difference in alpha power between conditions, where significant differences are indicated with ‘+’ markers (*p* < 0.05, *n* = 18, paired two-tailed Wilcoxon sign rank test, corrected for multiple comparisons over 96 electrodes by controlling the FDR at 0.05). During active and sham play, a significant decrease in alpha power was resolved over broad regions of the scalp, most notably over the left and right central electrodes. This suggests that the motor cortex was indeed engaged during sham play despite the absence of an overt motor task.

### Sham play elicits stronger late evoked responses over parietal cortex

Thus far, we pooled the data from all conditions in order to form a common set of evoked response components, and only evaluated differences in overall SRC. To find the origin of these differences, next we computed temporal and spatial response functions separately for each condition (see *Methods: Comparing spatial and temporal responses across conditions*). Active play should differ trivially from the inactive conditions due the evoked responses associated with motor and somatosensory activity. We therefore focused our analysis on the differences between sham play and passive viewing. The spatial and temporal patterns of evoked responses were largely preserved across conditions, except for the spatial pattern of the second component (Fig 4A-C, G-I). However, we observed significant differences in *magnitude* of the spatial and temporal responses. In particular, the spatial response was stronger during sham play compared to passive viewing, with a more negative response over the parietal cortex in the first component (Fig 4D). We also observed a significant increase in left central locations at the second component (Fig 4E). Furthermore, compared to passive viewing, the evoked responses measured during sham play were larger between 400 and 800 ms in components 1 and 2 (Fig 4G-H). This suggests that motor engagement may amplify late visual evoked responses that were generated downstream from the primary visual cortex.

**Fig. 4.**
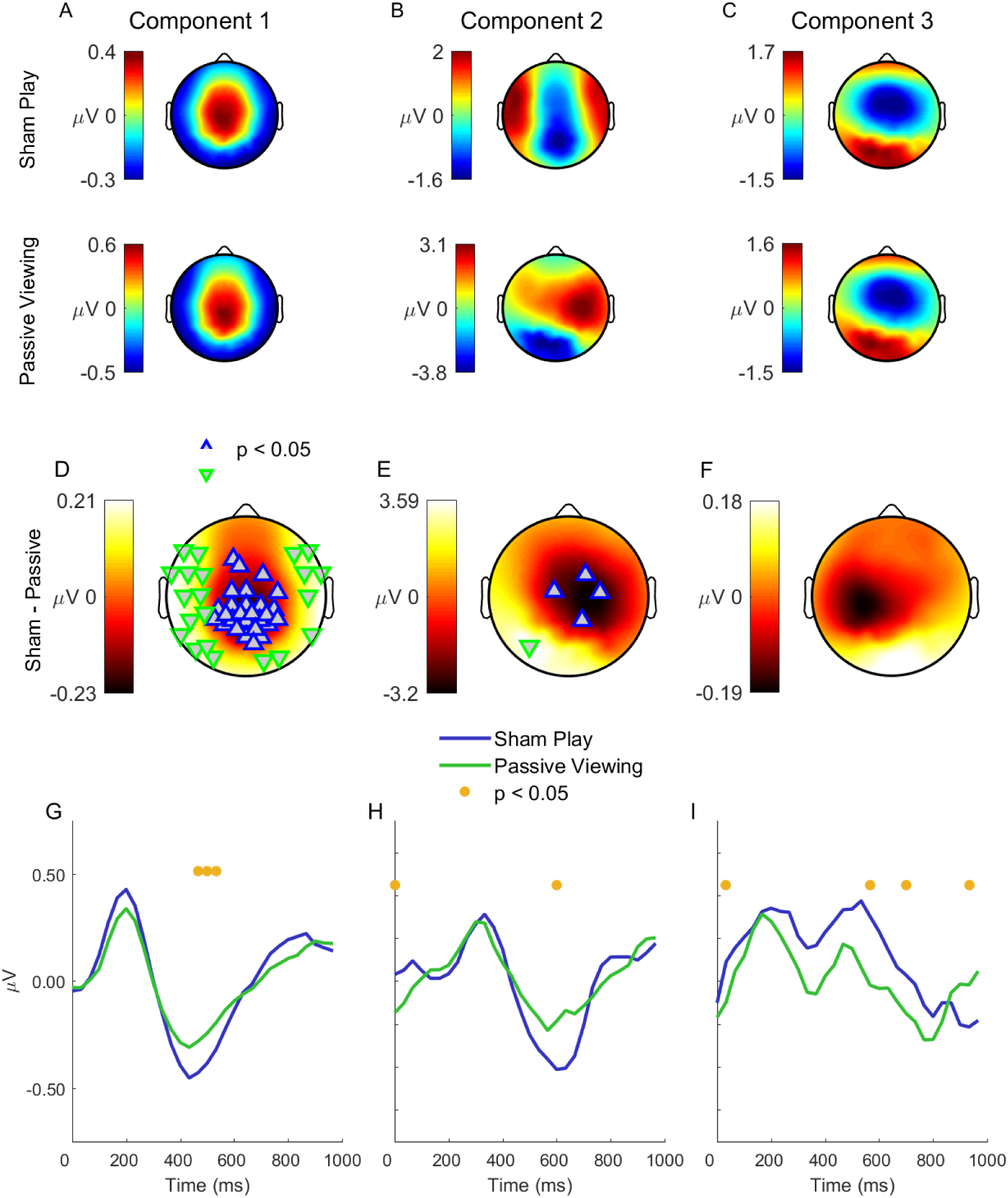
Sham play elicits stronger late evoked responses over parietal cortex. In order to compare the spatial and temporal characteristics of the evoked EEG responses, we performed the component analysis separately for each experimental condition. (**A**-**C**) Spatial response functions of the first three components, shown for sham play (top row) and passive viewing (bottom). (**D**-**F**) Sham play exhibited more negative responses over parietal cortex in the first component, while showing more positive responses at left central locations in the second component. The scalp maps indicate the spatial response difference between sham play and passive viewing, where significant effects are marked with triangles (corrected for multiple comparisons over 96 electrodes by controlling the FDR at 0.05, *n* = 18 subjects). (**G**-**I**) Temporal response functions of the first three components, shown separately for sham play and passive viewing. Time indicates the delay of the evoked EEG response relative to the stimulus presentation time. Sham play exhibited significantly stronger evoked potentials at late times (400-800 ms) in components 1 and 2. Shaded areas indicate standard error of the mean across subjects. Red dots indicate times exhibiting a significant difference (corrected for multiple comparisons over 30 time points by controlling the FDR at 0.05, *n* = 18 subjects).

### No SRC difference between counting task and passive viewing

One interpretation of the increased SRC during sham play is that participants paid more attention to the stimulus, thus enhancing visual evoked responses. To test this hypothesis, we repeated the study with a separate cohort of *n* = 20 subjects, but this time also asking subjects to view a prerecorded game while counting target items that appeared in the game (*n* = 66 items were presented during each race) – a task that required a high level of attention. This condition aimed to control for attention while removing any effects from engagement of the motor system.

In line with the initial study, we found increased SRC during both active and sham play compared to passive viewing (active vs passive: *z* = 2.61, *p* = 0.009; sham vs passive: *z* = 1.978, *p* = 0.048, *n* = 20, paired, two-tailed Wilcoxon signed rank test, Fig 5A). On the other hand, there was no significant difference in SRC between passive viewing and the counting task (*z* = 1.12, *p* = 0.26, *n* = 20). Although active and sham play elicited greater SRC than the counting task, the difference fell short of reaching significance (active vs count: *z* = 0.89, *p* = 0.37, sham vs count: *z* = 1.60, *p* = 0.11). The components measured during the follow-up study showed a strong resemblance to those found in the initial experiments (Fig S3).

**Fig. 5.**
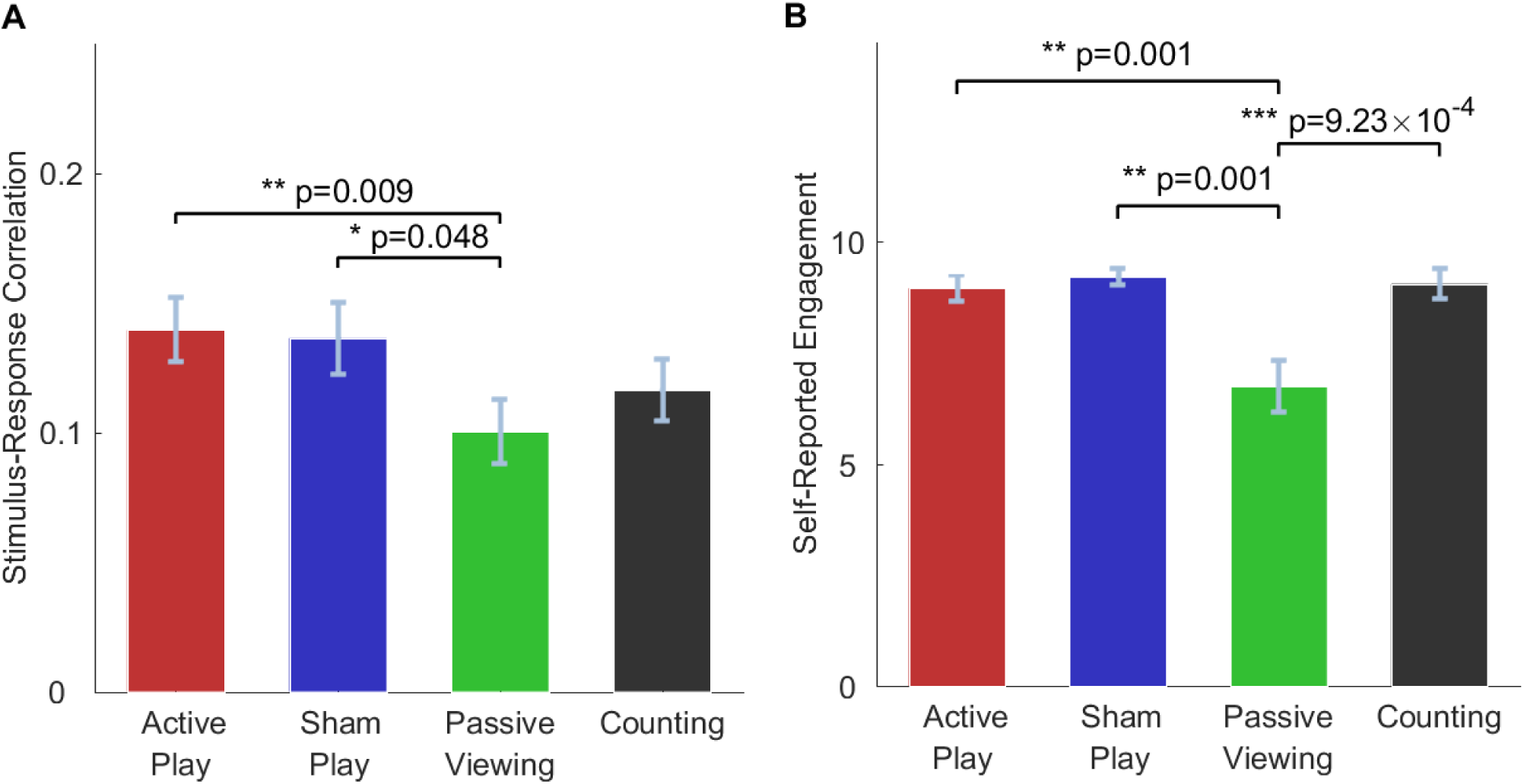
No enhancement in stimulus-response correlation during counting task. In order to control for the potential influence of top-down attention on the observed SRC increase, we performed a follow-up study including a condition where subjects were asked to count appearances of a target item on the screen while viewing playback of a pre-recorded game (“Counting”). (**A**) Reproducing the initial study, we observed a significant increase in SRC during both active and sham play relative to passive viewing (active play: *z* = 2.61, *p* = 0.009; sham play: *z* = 1.98, *p* = 0.048, *n* = 20, paired two-tailed Wilcoxon sign rank test). On the other hand, the counting task did not elicit a significant increase in SRC relative to passive viewing (*z* = 1.12, *p* = 0.26). Although sham play exhibited higher SRC compared to passive viewing, the difference fell short of significance (*z* = 1.60, *p* = 0.11). Bar heights depict the mean ± sem across *n* = 20 subjects. (**B**) Subjects reported significantly higher engagement during active play, sham play, and the counting task compared to passive viewing (*p* < 9.23 × 10^−4^, *n* = 20).

Self-reported engagement scores indicated that subjects were more engaged during active play, sham play, and the counting task relative to passive viewing (active play: *p* = 0.001; sham play: *p* = 0.001, counting task: *p* = 9.23 × 10^−4^, *n* = 20; Fig 5B). Of the 20 participants, only 6 subjects completed the experiment believing that their brain activity was controlling the video game.

Consistent with the findings of the initial study, we found reduced alpha power over the central electrodes during active and sham play relative to passive viewing (*p* < 0.05, *n* = 20, corrected for multiple comparisons by controlling the false discovery rate at 0.05; Fig S2A-B). On the other hand, there were no significant differences in alpha power between passive viewing and the counting task conditions (*p* > 0.05, *n* = 20; Fig S2A-B).

Finally, we probed differences in spatial and temporal response functions between sham play and the counting task (Fig 6). Interestingly, sham play exhibited a stronger (i.e., more negative) spatial response at occipital electrodes of the first three components (Fig 6D-F). Significant differences were also observed in a bilateral cluster of electrodes over frontotemporal cortex in component 3. Moreover, sham play also evoked stronger temporal responses at late delays (400 - 800 ms) in the first component (parietal cortex). Thus, the differences between sham play and counting mirrored those between sham play and passive viewing (Fig 4).

**Fig. 6.**
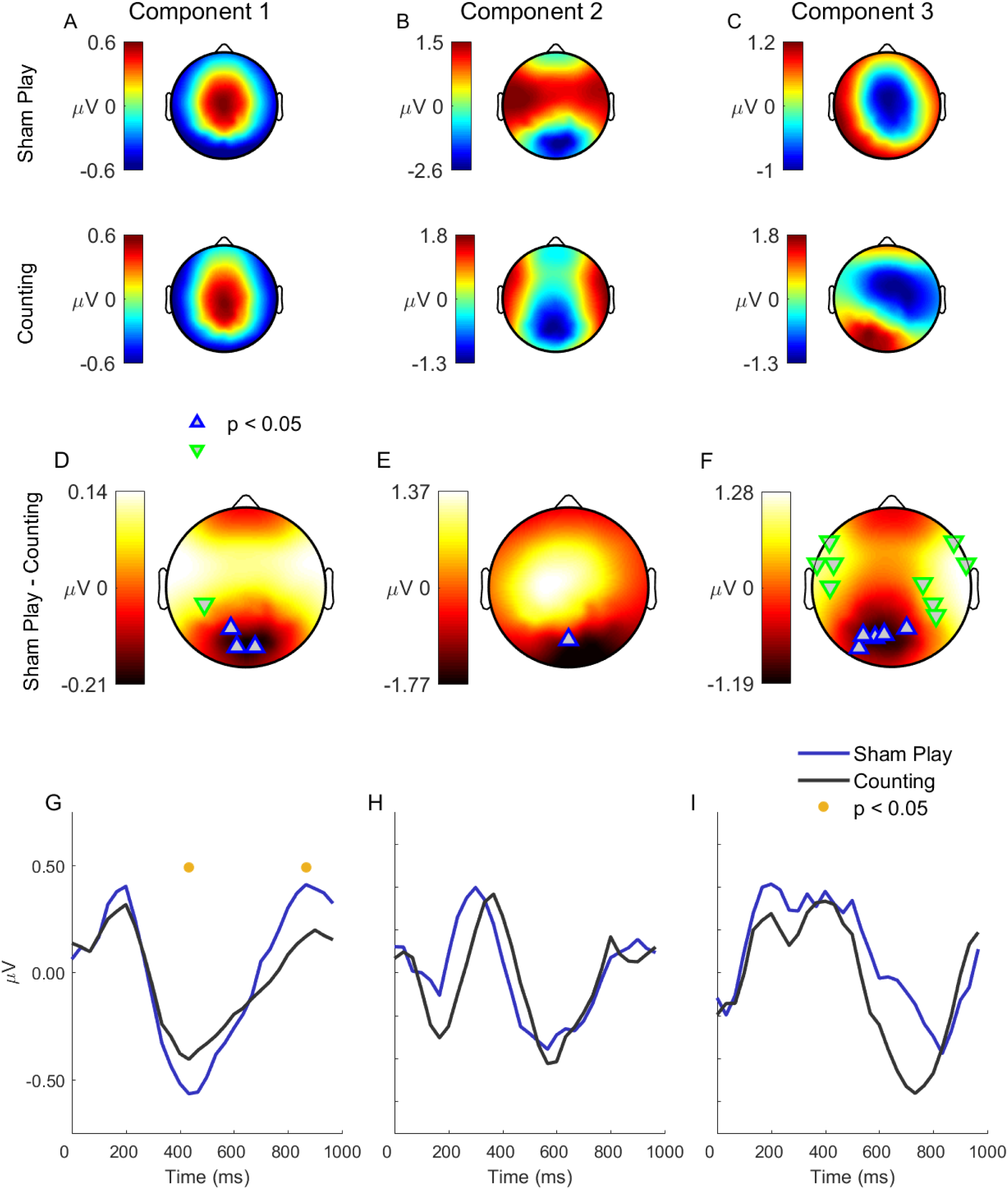
Visual evoked responses are stronger during sham play relative to counting. (**A**-**C**) To probe differences in the evoked responses between sham play and counting, we computed the spatial and temporal response functions separately for each condition. The spatial responses of the first three components are shown. (**D**-**F**) Sham play exhibited a stronger (i.e., more negative) spatial response at occipital electrodes of the first three components. Significant differences were also observed in a bilateral cluster of electrodes over frontotemporal cortex in component 3. (**G**-**I**) Temporal response functions of the first three components, shown separately for sham play and counting. Sham play evoked stronger temporal responses at late delays (400 - 800 ms) in the first component (centroparietal cortex). Compared with the difference between sham play and passive viewing (see Fig 4), these results suggest that the effect of attention on visual evoked responses differs qualitatively from the influence of motor engagement. Statistical analysis as in Figure 4

## Discussion

The findings of this study are consistent with the view that visual evoked responses are enhanced when vision guides motor control. The employment of a sham mitigated confounds from movement and somatosensation, while a reduction of alpha-band activity indicated that the motor cortex was indeed engaged despite the lack of overt actions. By asking subjects to count screen items during viewing, we attempted to control for increased attention in the play conditions relative to passive viewing. Indeed, sham play enhanced visual responses over passive viewing, but counting did not. However, the difference between sham play and counting fell short of reaching significance. Therefore, we cannot fully rule out that heightened attention contributed to the enhanced visual responses during sham play.

The main limitation of our study is thus that we were not able to fully disentangle the effects of an engaged motor system from that of increased top-down attention. The difficulty in deceiving subjects throughout this longer experiment (due to the addition of the fourth condition) likely contributed to this – only 6 of 20 subjects were fully deceived. It is likely that the visual evoked responses measured during sham play partially reflected brain states consistent with passive viewing. Note that despite this, we still found significant increases in the amplitudes of the spatial and temporal responses measured during sham play relative to counting (Fig 6). Another study limitation relates to possible differences in brain state between the sham play, passive viewing, and counting conditions that are separate from motor activation. For example, deceived subjects may have noticed discrepancies between the car’s movement and the intended command. The unpredictability of the stimulus in this case may have evoked stronger visual responses.

Visual stimuli containing objects that afford actions have been shown to increase visual spatial attention and amplify evoked responses, but only when the premotor and prefrontal cortices are activated (16, 26). This implies connectivity between premotor and prefrontal regions and the visual cortex, which has been shown anatomically in the primate brain (5–7). Here, the presence of the race car on the screen may have similarly amplified the evoked response to the optic flow stimulus. Note, however, that the modulation of visual responses required an active motor plan, in that the same actionable stimulus did not enhance visual responses during passive viewing.

Observing motor actions has been shown to generate imitative motor plans in the observer (27), but the role of these motor plans has been debated (28). One account is that the function of this motor activation is to generate a prediction of future perceptual input, thus bypassing the delays of sensory processing (29). During active and sham game play, study participants may have formed a prediction of the evolving optic flow stimulus, consistent with increased stimulus-driven activity over the central cortex (Fig 4). This interpretation of the results is consistent with the theory that perceived events and planned actions share a common representational domain (30).

In general, active and sham play may have exhibited stimulus-driven neural activity along a broader portion of the brain. For example, it is possible that the optic flow stimulus evoked correlated activity in dorsal regions downstream from striate visual cortex, such as the parietal or premotor cortices. Indeed, the strongest modulation of the evoked response, as well as alpha power, was seen over the parietal and central cortices (Figs 3-4). The first component was expressed over these areas (Fig 2). The posterior parietal cortex (PPC) has been shown to code motor intentions in the macaque (31), and it is tempting to speculate that a PPC-like component tracked the visual stimulus more faithfully in the sham play condition compared to the passive state. However, a limitation of our study is the poor spatial resolution of the scalp EEG, and the associated difficulties in recovering cortical sources from observed scalp topographies. The ill-posed nature of the EEG inverse problem is exacerbated when averaging scalp topographies over multiple subjects, as was implicitly done here. A natural extension of this work is thus to replicate the experiment with fMRI to glean insight into the brain areas driving the enhancement of visual evoked responses. However, note that the high temporal resolution of the EEG allowed us to measure fast evoked responses to the dynamic stimulus, which may not be feasible with fMRI due to the slowness of the hemodynamic response to neural activation.

The SRC approach taken here (22, 23) permitted the capture of continuous visual evoked responses during a sensorimotor task that more closely mimics real-world settings than conventional event-related designs that employ discrete stimuli. Moreover, we were able to capture several components of the neural response to the optic flow stimulus. Note that in our framework, the analogues of the classical visual event-related potential (ERP) are the temporal response functions shown in Fig 2A (second row), which are entirely analogous to what is extracted with multivariate regression techniques (24). These time courses indicate the brain’s response to an impulse of optic flow. In our framework, the response is expressed over a *set* of electrodes as depicted by the corresponding “spatial response function” (first row in Fig 2) (22). Note that while optic flow is a low-level feature of the visual stimulus, the neural response to it may be modulated by complex brain states such as anticipation, surprise, fear, or arousal. Thus, the neural activity that was measured here captured potentially more than the conventional visual evoked response. Note for example that the effects of an engaged motor cortex were to enhance late responses over central and parietal cortex. While not “visual” in the conventional sense, these evoked responses were nonetheless driven and thus correlated with the dynamic visual stimulus.

Regardless of the neural mechanism underlying the enhancement of visual processing during game play, our results provide an avenue for decoding active versus passive vision from non-invasive measurements of neural activity. By measuring the correlation between neural responses and a time-varying visual stimulus, one can extract an estimate of how active the viewer is. While here we measured SRC at the scale of a 3-minute trial, it can also be computed in finer time increments and tracked continuously. We speculate that there is a continuum between passive viewing and active control, and that the SRC can place the subjects onto this continuum on a moment-to-moment basis. In the future, wearable devices may be equipped with various sensors for capturing environmental stimuli in real-time (e.g. microphones and video cameras). Given the development of unobtrusive techniques for non-invasive sensing of neural activity (32), such as that from inside the ear canal (33), the SRC represents a natural technique for gleaning information about individual brain state in real-time. For example, it may be possible to decode spatial attention (34) by computing the SRC separately for multiple areas of the visual field or directions of incoming sound. There is already evidence that SRC can be used to determine speech comprehension in the context of hearing aids or capture a listener’s attention (23). An advantage of the SRC approach is that it is unsupervised, in that no learning procedure is required to, for example, learn patterns of neural activity that distinguish active from passive viewing.

Finally, an interesting facet of this work is that we were able to deceive a substantial number of our participants. In total, 19 of 38 participants completed the experiment with the belief that their brain activity was controlling game play during trials in which they actually viewed prerecorded stimuli. It is likely that the car racing video game employed in our study elicited stereotyped manual (and imagined) responses across subjects, thus contributing to the efficacy of deception. It is also notable that the sham play condition evoked strong neural activity over the parietal cortex (36), a region associated with visually guided movement planning and control. This suggests that such visuomotor pathways may be activated with only the perception of control. Aside of being an interesting psychological finding, this opens up new experimental paradigms for probing the brain under active scenarios.

## Methods

### Participants

All participants provided written informed consent in accordance with procedures approved by the Institutional Review Board of the City University of New York. In the initial study, 18 healthy human subjects (9 females) aged 20±1.56 participated. For the follow-up experiment testing the effects of attention, we recruited a new cohort of 24 healthy subjects (14 females, 20 ± 3.01 years).

### Video game stimulus

We employed the open-source car racing game SuperTuxKart, in which participants navigate vehicles around a track against simulated opponents. All experimental trials were conducted on the default course and spanned three laps in “easy” mode. The average trial had a duration of 175.9±5.51 seconds. In the initial study, we removed several graphical items from the stimulus such that the video stimulus consisted of only the race car, track, and opponents. To generate the stimuli employed during the sham play and passive viewing conditions, we recorded several races for subsequent playback during the experiments. A non-participant played 4 races, with 2 serving as stimuli during the sham play condition and the other 2 employed during the passive viewing condition. With the exception of active play, which produces unique stimuli during each trial, all subjects experienced the same stimuli.

The stimulus was presented on a high-definition Dell 24-inch UltraSharp Monitor (1920-by-1080 pixels) at a frame rate of 60 Hz. Subjects viewed the stimulus in a dark room at a viewing distance of 60 cm. The game’s sound was muted during the experiment. The video frame sequence of each race was captured with the open-source Open Broadcaster Software at the native resolution and frame rate. In order to subsequently synchronize the video frame sequence with the recordings of the EEG, a 30-by-30 pixel square was flashed in the top right corner of the display throughout each trial. These were not visible to the subject, but a photodiode registered these markers and transmitted an electrical pulse to the EEG recorder with low latency.

### Experimental procedures

Initial study participants experienced two trials of the video game stimulus in each of 3 conditions: “active play”, “sham play”, and “passive viewing”. The ordering of the conditions was randomized and counter-balanced across subjects. Subjects were permitted one practice trial of the video game prior to commencing the experiment. During active play, subjects controlled the game via keyboard presses made with the right hand: the left and right keys controlled steering, while the up and down keys produced acceleration and braking, respectively. Prior to the first sham play trial, subjects were falsely told that their brain activity will be controlling the video game, and that they should imagine the intended command. Moreover, we primed the participants by implementing a mock calibration of a brain computer interface before playback of the sham play races, during which subjects were asked to imagine game controls (accelerate, brake, steer left, steer right). During passive viewing trials, subjects were instructed to freely view playback of a previously recorded game. Upon completion of the experiment, participants filled out a survey reporting their experienced “engagement” during each condition. Scores ranged from 1 (“not engaged”) to 10 (“fully engaged”). Following the survey, subjects were informed of the deception task, and were asked whether they had become aware of the fact that their brain activity was not controlling game play.

In the follow-up experiment, we repeated the same three conditions (active play, sham play and passive viewing) and included a fourth “counting” condition. In this condition, subjects were instructed to view pre-recorded playback of the game while simultaneously counting the total number of appearances of a target item (i.e., a gift-box) on the track. Upon completion of each trial, subjects were asked to recall the total number of times that the item appeared. The correct number of items was 66, with items appearing regularly over the approximately three-minute trial. As in the initial experiment, the ordering of the conditions was randomized and counterbalanced across subjects, and each condition of the game was repeated twice.

### EEG acquisition and preprocessing

The scalp electroen-cephalogram (EEG) was acquired with a 96-electrode cap (custom montage with dense coverage of the occipital region) housing active electrodes connected to a Brain Products ActiChamp system and Brain Products DC Amplifier (Brain Vision GmbH, Munich, Germany). The EEG was sampled at 500 Hz, digitized with 24 bits per sample, and transmitted to a recording computer via the Lab Streaming Layer software (37), which ensured precise temporal alignment between the EEG and video frame sequence.

EEG data was imported into the Matlab software (Mathworks, Natick, MA) and analyzed with custom scripts. Data was downsampled to 30 Hz in accordance with the Nyquist rate afforded by the 60 Hz frame rate, followed by high-pass filtering at 1 Hz to remove slow drifts. To remove gross artifacts from the data, we employed the robust PCA technique (38), which provides a low-rank approximation to the data and thereby removes sparse noise from the recordings. Note that due to volume conduction, sparse EEG components are generally artifacts. We employed the robust PCA implementation of Lin et al. (39) with the default hyperparameter of *λ* = 0.5. To reduce the contamination of EEG from eye movements, we linearly regressed out the activity of four “virtual” electrodes constructed via summation or subtraction of appropriately selected frontal electrodes. These virtual electrodes were formed to strongly capture the activity produced by eye blinks and saccades. To further denoise the EEG, we rejected electrodes whose mean power exceeded the mean of all channel powers by four standard deviations. Within each channel, we also rejected time samples (and its adjacent samples) whose amplitude exceeded the mean sample amplitude by four standard deviations. We repeated the channel and sample rejection procedures over three iterations.

During the follow-up experiment, we experienced problems with synchronizing the EEG recordings with the video frame sequence. This was diagnosed by visually inspecting the temporal alignment between the periodic flashes registered by the recorder’s auxiliary channel and the corresponding event in the recorded video frame sequence. In 4 of 24 subjects, we observed misalignment exceeding 300 ms and excluded these 4 subjects from all analyses.

### Stimulus feature extraction

Video frames were downsampled to a resolution of 320-by-180 pixels to reduce data size, and then converted to grayscale images. Optical flow was computed with the Horn-Schunk method as implemented in the MATLAB Computer Vision System Toolbox (40). For each frame, we computed the mean (across pixels) of the magnitude of the optical flow vector. Temporal contrast was constructed by taking the mean (across pixels) of the frame-to-frame difference of the video sequence (22). The resulting time series were z-scored prior to SRC analysis.

### Stimulus-response correlation (SRC)

To measure the correlation between the time-varying stimulus feature *s*(*t*) and the *D* dimensional evoked neural response *r*_*i*_(*t*), *i* ∈ 1, 2, …, *D*, we employed the multidimensional SRC technique developed in Dmochowski et al. (22). This regression approach leveraged Canonical Correlation Analysis and was independently developed in de Cheveigné et al. (23). The approach consists of temporally filtering the stimulus:

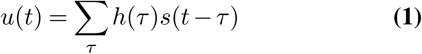

and spatially filtering the neural response:

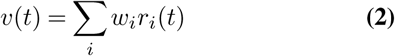

to produce stimulus component *u*(*t*) and response component *v*(*t*) that exhibit maximal correlation:

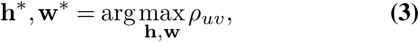

where **h**^*^ = [*h*(1) … *h*(*L*)]^*T*^ are the optimal temporal filter coefficients of the *L*-length filter and **w**^*^ = [*w*_1_ … *w*_*D*_]^*T*^ are the optimal spatial filter coefficients, and where *ρ*_*uv*_ is the Pearson correlation coefficient between *u*(*t*) and *v*(*t*). The solution to (3) is given by Canonical Correlation Analysis (41) and consists of pairs of projection vectors 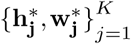 that yield a set of maximally correlated components *u*_*j*_(*t*) and *v*_*j*_(*t*) with corresponding correlation coefficients that decrease in magnitude 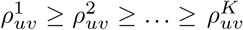. Note here that we regularized the CCA solution by truncating the eigenvalue spectrum of the EEG covariance matrix to *K* = 11 dimensions, as this value explained over 99% of the variance in the data. Encompassing all components, the total correlation between the stimulus and response is given by:

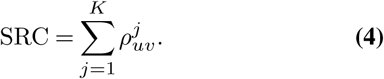

With the exception of the results presented in Figures 4 and 6, the CCA filters were computed after pooling data from all conditions. In this manner, SRCs were computed over a common basis for all conditions.

The filter coefficients *h*_*j*_(*t*) are equivalent to the “temporal response function” extracted with conventional multivariate regression (24). Note that here *j* represents a specific component, rather than a specific electrode as in Crosse et al. (24). One can also conceive of *h*_*j*_(*t*) as the temporal response function of a virtual electrode or “source” *j*, extracted from the EEG with the spatial filter **w**_*j*_. The corresponding “forward model”, **a**_*j*_, can be obtained following conventional approaches in EEG (42, 43). In Dmochowski et al. (22) we show that this forward model **a**_*j*_ is the equivalent of a “spatial response function”, in that the total evoked response is the product of the spatial and temporal responses, summed over all components: 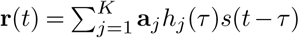.

### Alpha power analysis

To test for differences in alpha power between conditions (Figs 3, S2), we temporally filtered the EEG response of each electrode *r*_*i*_(*t*), *i* = 1, …, *D*, to the alpha band (8-12 Hz) using a fourth order Butterworth filter. We then measured the alpha power at each electrode by computing the temporal mean square of the filter output. Alpha power was averaged across the two trials performed by each participant prior to statistical tests.

### Statistical testing

We tested for conditional differences in SRC, self-reported engagement, and alpha power by conducting paired, two-tailed Wilcoxon signed-rank tests on sets of *n* = 18 (or *n* = 20 for the follow-up study) samples in each condition, with each exemplar corresponding to a subject.

### Comparing spatial and temporal responses across conditions

To test for spatial and temporal differences in the visual evoked responses across conditions (Fig 4), we computed response functions separately for active play, sham play, and passive viewing, and counting. In order to obtain the conditional spatial response functions, we first filtered the stimulus with the first three temporal response functions (shown in Fig 2A, second row). This yielded three distinct (filtered) versions of the optic flow, *u*_*j*_(*t*), *j* = 1 … 3. We then performed a multivariate spatial regression from the scalp EEG onto this filtered optic flow, but separately for each condition. The result represents the spatial distribution of the EEG response **a**_*j*_ to the filtered optic flow stimulus in each condition (Fig 4A-C). Analogously, to obtain the temporal response function for each condition, we spatially filtered the EEG with the first three CCA-derived filters. This is equivalent to generating three virtual electrodes *v*_*j*_(*t*), *j* = 1 … 3. We then performed a temporal regression from the optic flow stimulus onto these spatially filtered neural responses. The resulting time courses represent the dynamics *h*_*j*_(*t*) of the visual evoked response in each condition (Fig 4G-I).

To test for significant differences between conditions (sham versus passive: Fig 4; sham play versus counting Fig 6), we computed the difference of the values **a**_*j*_ (or *h*_*j*_(*t*)) between passive and sham conditions (Fig 4D-F). These differences were measured on group-averaged spatial response or temporal response functions. Statistical significance was conducted with a permutation test. A null distribution of conditional differences was generated by randomly swapping subject assignment between sham play and passive viewing (without replacement) over 1000 random assignments. To correct for multiple comparisons, we controlled the false discovery rate at 0.05 across the 96 electrodes and 30 times points, respectively.

---

**Fig. S1.**
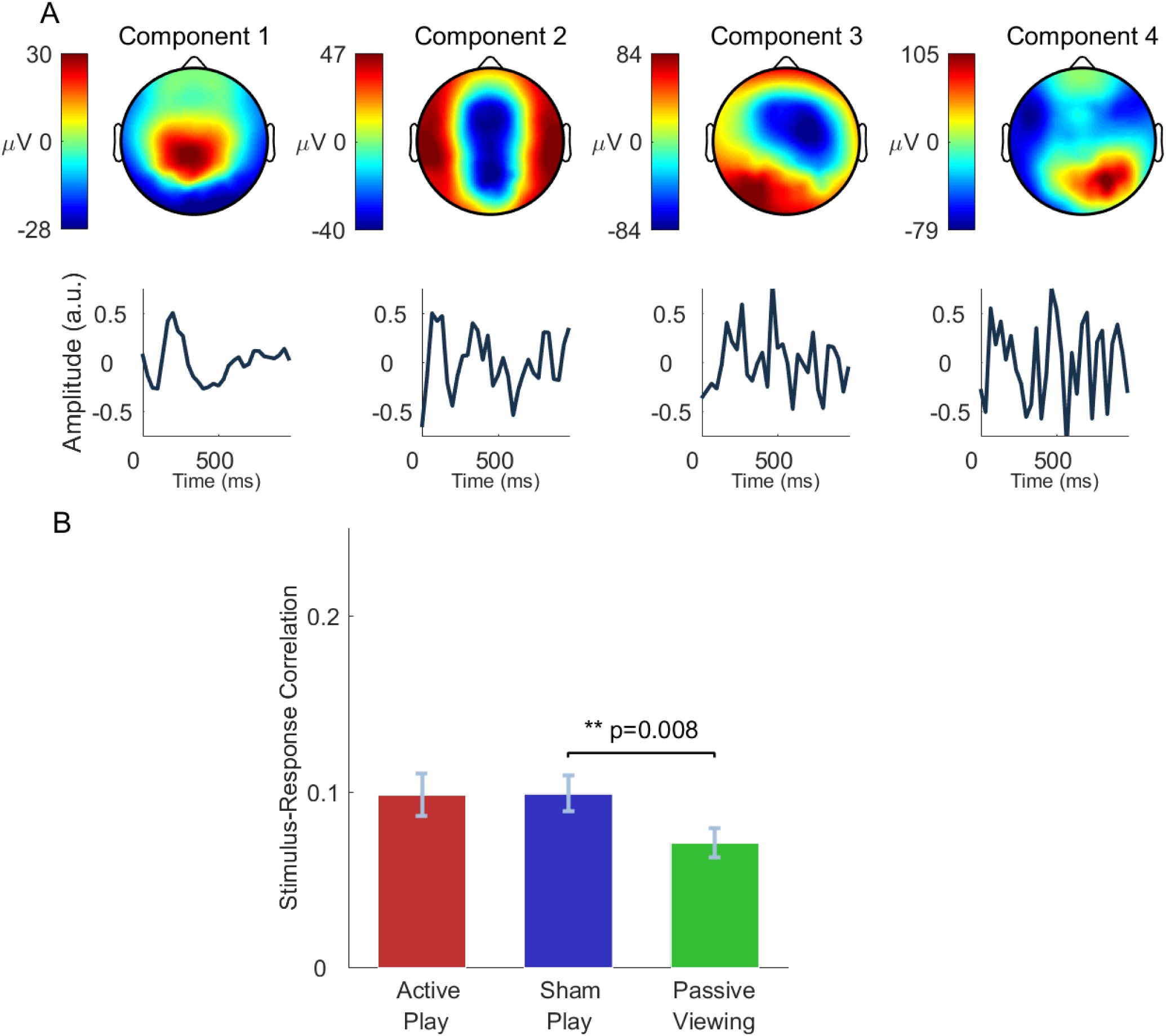
Reproducibility of effect with temporal visual contrast. We tested whether the effect of motor engagement on visual evoked responses would be reproduced when regressing the EEG onto visual contrast, as opposed to the optic flow used in the main analysis; compare with Fig 2. (**A**) (Top row) The spatial response functions of the first four components are largely consistent with those found using optic flow. (Bottom row) The temporal response function of the first component is consistent with that found using optic flow. The remaining components exhibit more high-frequency activity than found in the main analysis. (**B**) Reproducing the effect found with optic flow, sham play yielded significantly higher total SRC compared to passive viewing (*p* = 0.008, two-tailed, paired Wilcoxon signed rank test, *n* = 18). Active play also produced higher SRC compared to passive viewing, but the difference fell short of statistical significance (*p* = 0.14).

**Fig. S2.**
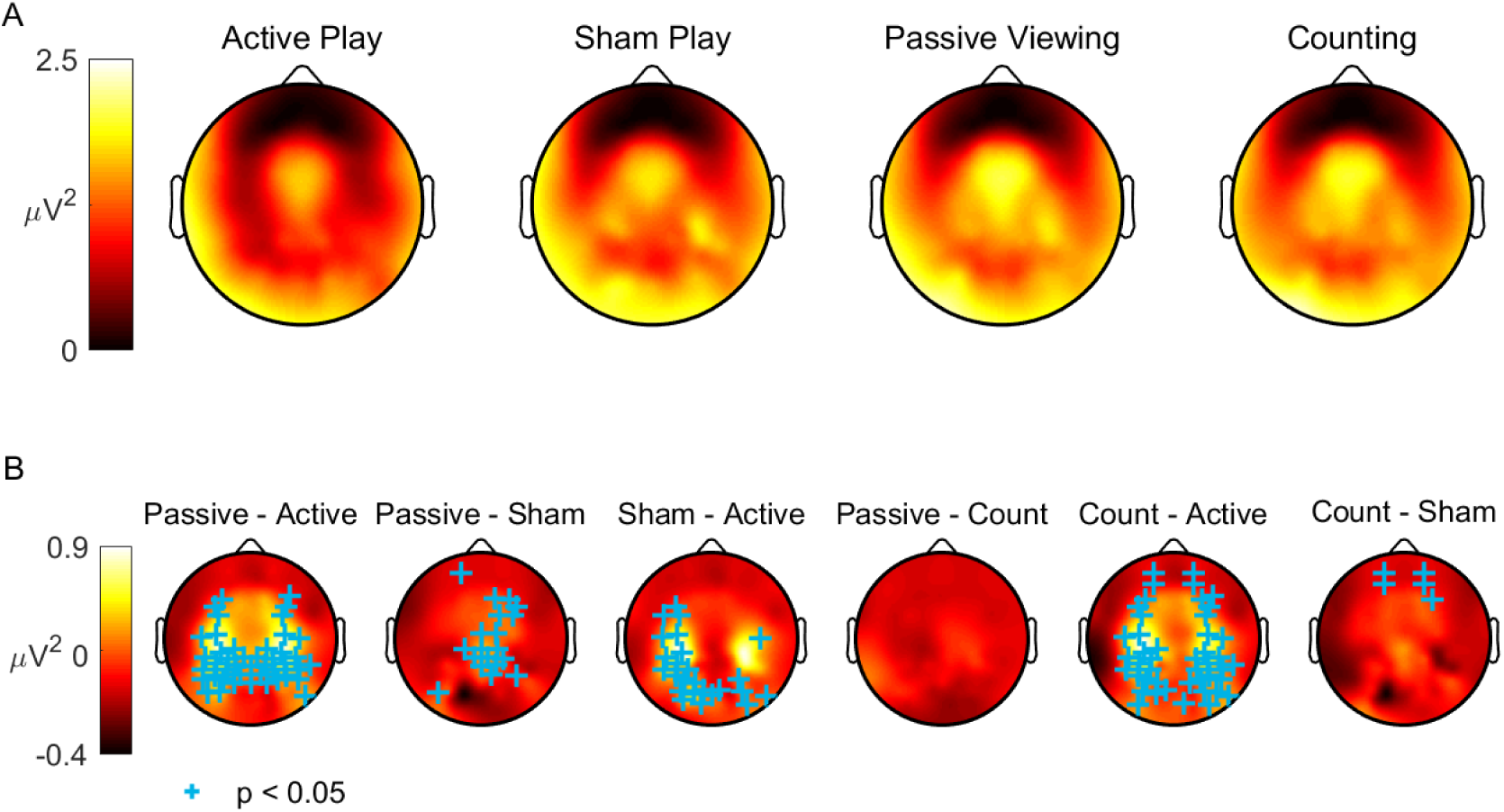
No difference in alpha power between counting and passive viewing. In the initial study, we found a significant decrease in alpha power during sham play relative to passive viewing, suggestive of motor cortex recruitment during sham play. (**A**) We measured alpha power for all four conditions in the follow-up study: active play, sham play, passive Viewing, and counting. (**B**) Consistent with the findings of the initial study, we found reduced alpha power over central and parietal electrodes during active and sham play relative to passive viewing (*p* < 0.05, *n* = 20, corrected for multiple comparisons by controlling the false discovery rate at 0.05). On the other hand, there were no significant differences in alpha power between passive viewing and the counting task (*p* > 0.05, *n* = 20).

**Fig. S3.**
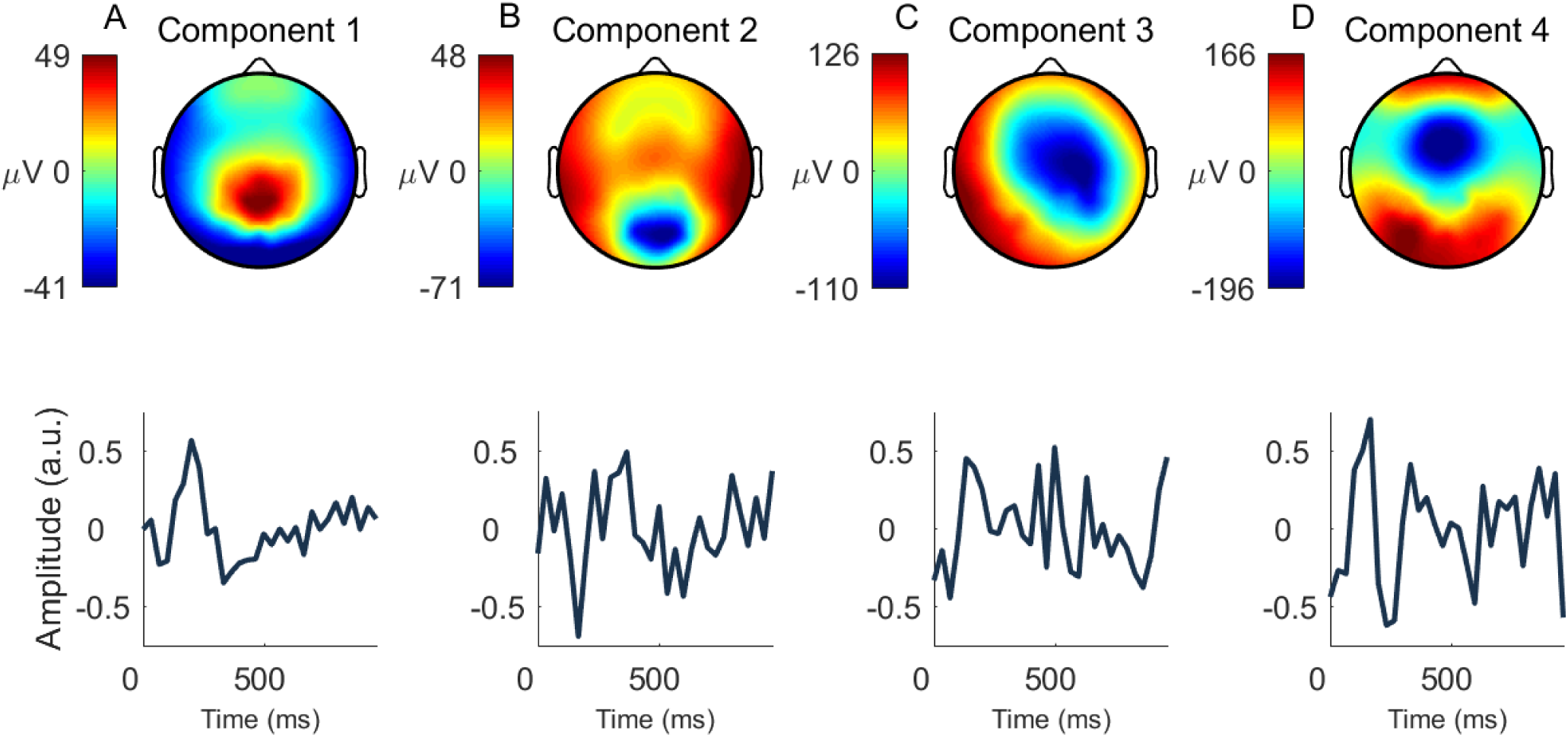
Evoked response patterns are reproduced in follow up study. (Top row) Spatial response components of the first four components computed using optic flow bear a strong resemblance to those found in the initial study (compare with Fig 2A). The strongest component is expressed over centroparietal electrodes. (Bottom row) The corresponding temporal response functions represent the time course of the evoked response to the optic flow stimulus.

## References

1. Mortimer Mishkin, Leslie G Ungerleider, and Kathleen A Macko. Object vision and spatial vision: two cortical pathways. Trends in neurosciences, 6:414–417, 1983.

2. AD Milner and MA Goodale. Two visual systems re-viewed. Neuropsychologia, 2008.

3. Melvyn A Goodale and A David Milner. Separate visual pathways for perception and action. Trends in neurosciences, 15(1):20–25, 1992.

4. David Milner and Mel Goodale. The visual brain in action. Oxford University Press, 2006.

5. Malcolm P Young. The organization of neural systems in the primate cerebral cortex. Proceedings of the Royal Society of London. Series B: Biological Sciences, 252(1333):13–18, 1993.

6. Steven P Wise, Driss Boussaoud, Paul B Johnson, and Roberto Caminiti. Premotor and parietal cortex: corticocortical connectivity and combinatorial computations. Annual review of neuroscience, 20(1):25–42, 1997.

7. David C Van Essen. Corticocortical and thalamocortical information flow in the primate visual system. Progress in brain research, 149:173–185, 2005.

8. Antonino Casile and Martin A Giese. Nonvisual motor training influences biological motion perception. Current Biology, 16(1):69–74, 2006.

9. Heiko Hecht, Stefan Vogt, and Wolfgang Prinz. Motor learning enhances perceptual judgment: A case for action-perception transfer. Psychological research, 65(1):3–14, 2001.

10. Annerose Engel, Michael Burke, Katja Fiehler, Siegfried Bien, and Frank Rösler. Motor learning affects visual movement perception. European Journal of Neuroscience, 27(9): 2294–2302, 2008.

11. Andreas Wohlschläger. Visual motion priming by invisible actions. Vision research, 40(8): 925–930, 2000.

12. Liana E Brown, Elizabeth T Wilson, Melvyn A Goodale, and Paul L Gribble. Motor force field learning influences visual processing of target motion. Journal of Neuroscience, 27 (37):9975–9983, 2007.

13. Hannah Barbara Helbig, Jasmin Steinwender, Markus Graf, and Markus Kiefer. Action observation can prime visual object recognition. Experimental brain research, 200(3-4): 251–258, 2010.

14. Fani Loula, Sapna Prasad, Kent Harber, and Maggie Shiffrar. Recognizing people from their movement. Journal of Experimental Psychology: Human Perception and Performance, 31 (1):210, 2005.

15. Günther Knoblich and Rüdiger Flach. Predicting the effects of actions: Interactions of perception and action. Psychological science, 12(6):467–472, 2001.

16. Todd C Handy, Scott T Grafton, Neha M Shroff, Sarah Ketay, and Michael S Gazzaniga. Graspable objects grab attention when the potential for action is recognized. Nature neuroscience, 6(4):421, 2003.

17. Maha Adamo and Susanne Ferber. A picture says more than a thousand words: Behavioural and erp evidence for attentional enhancements due to action affordances. Neuropsychologia, 47(6):1600–1608, 2009.

18. Agnieszka Wykowska and Anna Schubö. Action intentions modulate allocation of visual attention: electrophysiological evidence. Frontiers in Psychology, 3:379, 2012.

19. Heath Matheson, Aaron J Newman, Jason Satel, and Patricia McMullen. Handles of manipulable objects attract covert visual attention: Erp evidence. Brain and cognition, 86:17–23, 2014.

20. Giuseppe di Pellegrino, Robert Rafal, and Steven P Tipper. Implicitly evoked actions modulate visual selection: Evidence from parietal extinction. Current Biology, 15(16):1469–1472, 2005.

21. Liyu Cao and Barbara Händel. Walking enhances peripheral visual processing in humans. PLoS biology, 17(10):e3000511, 2019.

22. Jacek P Dmochowski, Jason J Ki, Paul DeGuzman, Paul Sajda, and Lucas C Parra. Extracting multidimensional stimulus-response correlations using hybrid encoding-decoding of neural activity. NeuroImage, 2017.

23. Alain de Cheveigné, Daniel DE Wong, Giovanni M Di Liberto, Jens Hjortkjaer, Malcolm Slaney, and Edmund Lalor. Decoding the auditory brain with canonical component analysis. NeuroImage, 172:206–216, 2018.

24. Michael J Crosse, Giovanni M Di Liberto, Adam Bednar, and Edmund C Lalor. The multivariate temporal response function (mtrf) toolbox: a matlab toolbox for relating neural signals to continuous stimuli. Frontiers in human neuroscience, 10:604, 2016.

25. Jaime A Pineda. The functional significance of mu rhythms: translating “seeing” and “hearing” into “doing”. Brain research reviews, 50(1):57–68, 2005.

26. Glyn W Humphreys, Eun Young Yoon, Sanjay Kumar, Vaia Lestou, Keiko Kitadono, Katherine L Roberts, and M Jane Riddoch. The interaction of attention and action: From seeing action to acting on perception. British Journal of Psychology, 101(2):185–206, 2010.

27. Giacomo Rizzolatti and Laila Craighero. The mirror-neuron system. Annu. Rev. Neurosci., 27:169–192, 2004.

28. Giacomo Rizzolatti, Luciano Fadiga, Vittorio Gallese, and Leonardo Fogassi. Premotor cortex and the recognition of motor actions. Cognitive brain research, 3(2):131–141, 1996.

29. Margaret Wilson and Günther Knoblich. The case for motor involvement in perceiving con-specifics. Psychological bulletin, 131(3):460, 2005.

30. Wolfgang Prinz. Perception and action planning. European journal of cognitive psychology, 9(2):129–154, 1997.

31. James W Gnadt and Richard A Andersen. Memory related motor planning activity in posterior parietal cortex of macaque. Experimental brain research, 70(1):216–220, 1988.

32. Alexander J Casson, David C Yates, Shelagh JM Smith, John S Duncan, and Esther Rodriguez-Villegas. Wearable electroencephalography. IEEE engineering in medicine and biology magazine, 29(3):44–56, 2010.

33. David Looney, Preben Kidmose, Cheolsoo Park, Michael Ungstrup, Mike Lind Rank, Karin Rosenkranz, and Danilo P Mandic. The in-the-ear recording concept: User-centered and wearable brain monitoring. IEEE pulse, 3(6):32–42, 2012.

34. Gi-Yeul Bae and Steven J Luck. Dissociable decoding of spatial attention and working memory from eeg oscillations and sustained potentials. Journal of Neuroscience, 38(2): 409–422, 2018.

35. Ivan Vladimirov Iotzov and Lucas C Parra. Eeg can predict speech intelligibility. Journal of Neural Engineering, 2019.

36. Christopher A Buneo and Richard A Andersen. The posterior parietal cortex: sensorimotor interface for the planning and online control of visually guided movements. Neuropsychologia, 44(13):2594–2606, 2006.

37. C Kothe. Lab streaming layer (lsl). https://github.com/sccn/labstreaminglayer. Accessed on October, 26:2015, 2014.

38. Emmanuel J Candès, Xiaodong Li, Yi Ma, and John Wright. Robust principal component analysis? Journal of the ACM (JACM), 58(3):11, 2011.

39. Zhouchen Lin, Minming Chen, and Yi Ma. The augmented lagrange multiplier method for exact recovery of corrupted low-rank matrices. arXiv preprint 1009.5055, 2010.

40. Berthold KP Horn and Brian G Schunck. Determining optical flow. Artificial intelligence, 17 (1-3):185–203, 1981.

41. Harold Hotelling. Relations between two sets of variates. Biometrika, 28(3/4):321–377, 1936.

42. Stefan Haufe, Frank Meinecke, Kai Görgen, Sven Dähne, John-Dylan Haynes, Benjamin Blankertz, and Felix Bießmann. On the interpretation of weight vectors of linear models in multivariate neuroimaging. Neuroimage, 87:96–110, 2014.

43. Lucas C Parra, Clay D Spence, Adam D Gerson, and Paul Sajda. Recipes for the linear analysis of eeg. Neuroimage, 28(2):326–341, 2005.

